# Tributyltin induces a transcriptional response without a brite adipocyte signature in adipocyte models

**DOI:** 10.1101/328203

**Authors:** Stephanie Kim, Amy Li, Stefano Monti, Jennifer J. Schlezinger

## Abstract

Tributyltin (TBT), a peroxisome proliferator-activated receptor γ (PPARγ)/retinoid X receptor (RXR) ligand and founding member of the environmental obesogen chemical class, induces adipocyte differentiation and suppresses bone formation. A growing number of environmental PPARγ ligands are being identified. However, the potential for environmental PPARγ ligands to induce adverse metabolic effects has been questioned because PPARγ is a therapeutic target in treatment of type II diabetes. We evaluated the molecular consequences of TBT exposure during bone marrow multipotent mesenchymal stromal cell (BM-MSC) differentiation in comparison to rosiglitazone, a therapeutic PPARγ ligand, and LG100268, a synthetic RXR ligand. Mouse primary BM-MSCs (female, C57BL/6J) undergoing bone differentiation were exposed to maximally efficacious and human relevant concentrations of rosiglitazone (100 nM), LG100268 (100 nM) or TBT (80 nM) for 4 days. Gene expression was assessed using microarrays, and *in silico* functional annotation was performed using pathway enrichment analysis approaches. Pathways related to osteogenesis were downregulated by all three ligands, while pathways related to adipogenesis were upregulated by rosiglitazone and TBT. However, pathways related to mitochondrial biogenesis and <u>br</u>own-in-wh<u>ite</u> (brite) adipocyte differentiation were more significantly upregulated in rosiglitazone-treated than TBT-treated cells. The lack of induction of genes involved in adipocyte energy dissipation by TBT was confirmed by an independent gene expression analysis in BM-MSCs undergoing adipocyte differentiation and by analysis of a publically available 3T3 L1 data set. Furthermore, rosiglitazone, but not TBT, induced mitochondrial biogenesis. This study is the first to show that an environmental PPARγ ligand has a limited capacity to induce health promoting activities of PPARγ.

## Introduction

Metabolic disruptors are chemicals that play a role in altering susceptibility to obesity, metabolic syndrome and related metabolic disorders including type 2 diabetes (Heindel, et al., 2015). Identification of this new class of chemicals evolved from the initial recognition that environmental chemicals (originally termed environmental obesogens) play a role in the metabolic programming of obesity risk following *in utero* exposure (Grün and Blumberg 2006).

The hunt for metabolism disrupting chemicals has focused on identifying ligands of peroxisome proliferator activated receptor γ (PPARγ), a nuclear receptor that is essential for white, <u>br</u>own-in-wh<u>ite</u> (brite) and brown adipocyte differentiation and mature adipocyte, maintenance, function and survival (Anghel et al. 2007; Imai et al. 2004; Tontonoz et al. 1994). PPARγ is a ligand activated transcription factor, and its activation regulates energy homeostasis by both stimulating storage of excess energy as lipids in white adipocytes and stimulating energy utilization by triggering mitochondrial biogenesis, fatty acid oxidation and thermogenesis in brite adipocytes. PPARγ forms a heterodimeric complex with retinoid X receptors (RXR) and binds to the consensus response element (5’-CAAAACAGGTCANAGGTCA-3’) (Juge-Aubry et al. 1997). Structurally diverse ligands can bind to PPARγ, because of its large binding pocket, including full agonist, partial agonist and antagonist ligand types (Garcia-Vallvé et al. 2015; Nolte et al. 1998). However, not all ligands interact with the binding pocket of PPARγ in the same manner. For instance, greater stabilization of helix 12 by full agonists results in their efficacious transcriptional activation of PPARγ compared to partial agonists (Pochetti et al. 2007).

The organotin tributyltin (TBT), which inspired the coining of the term environmental obesogen, is an environmental PPARγ ligand (Grün et al. 2006; Kanayama et al. 2005). TBT is a highly potent PPARγ ligand, which induces differentiation of bone marrow and adipose-derived multipotent stromal cells (MSCs) into adipocytes and suppresses bone formation (Baker et al. 2015; Grün et al. 2006; Li et al. 2011; Watt and Schlezinger 2015; Yanik et al. 2011). Prenatal TBT exposure in mice leads to epigenetic changes in mesenchymal stem cells that favor adipogenesis at the expense of osteogenesis (Chamorro-García et al. 2013), reduced ossification of the skeleton (Tsukamoto et al. 2004), and increased adiposity in adulthood (Grün et al. 2006). In adults, TBT not only induces weight gain but also metabolic disruption (i.e. hepatic steatosis, hyperinsulinemia and hyperleptinemia) (Bertuloso et al. 2015; Zuo et al. 2011) and adipocyte formation in bone (Baker et al. 2017). Since the discovery of TBT’s ability to activate PPARγ, a growing number of environmental PPARγ ligands that stimulate adipogenesis have been identified (e.g. organotins, phthalates, parabens, triflumizole, tetrabromobisphenol A, triphenyl phosphate) (Hu et al. 2013; Kanayama et al. 2005; Li et al. 2012; Pillai et al. 2014; Riu et al. 2011; Watt and Schlezinger 2015).

Moreover, TBT is not only a PPARγ agonist, but also is an agonist for retinoid X receptors (RXR) α and β (le Maire et al. 2009; Shiizaki et al. 2014). The dual ligand nature of the organotins raises the possibility that RXR could contribute to TBT-induced effects on MSC differentiation, independently of PPARγ. RXRs heterodimerize with and are essential for transcriptional activation of multiple nuclear receptors; further, they also act as homodimers (Kojetin et al. 2015). Permissive RXR heterodimer partners may be activated by ligand binding to either the partner or to RXR (Lammi et al. 2008; Schulman et al. 1998). In reporter assays, TBT was shown to activate liver X receptor (LXR), nuclear receptor related 1 (NURR1), PPARγ and PPARδ, which are permissive RXR heterodimeric partners (Grün et al. 2006). We have previously shown that TBT can act through RXR homodimers and RXR/PPARγ heterodimers to induce adipogenesis and suppress osteogenesis in mouse bone marrow MSCs, as well as can induce LXR-dependent gene transcription (Baker et al. 2015).

PPARγ is a therapeutic target in the treatment of type II diabetes. The PPARγ ligands Rosiglitazone (Avandia™) and pioglitazone (Actos™) are prescribed to increase insulin sensitivity and reduce blood glucose (Nolte et al. 1998). This has raised the question as to whether environmental PPARγ ligands are simply adipogenic or will induce adverse metabolic effects. To begin to address this question, we compared the TBT-induced transcriptional response in bone marrow MSCs (BM-MSCs) to those induced by rosiglitazone and LG100268, a synthetic RXR agonist. Bioinformatics analyses of our BM-MSC-derived data and of a publically available, independently generated 3T3 L1 dataset revealed that while TBT is potent and efficacious at stimulating white adipogenesis and lipid accumulation, it does not efficiently activate energy dissipating pathways (e.g. mitochondrial biogenesis, adipocyte browning).

## Methods

### Chemicals

DMSO was purchased from American Bioanalytical (Natick, MA). Rosiglitazone (Rosi) was from Cayman Chemical (Ann Arbor, MI). Human insulin, dexamethasone, 3-isobutyl-1-methylxanthine (IBMX), LG100268 (LG268), and tributyltin (TBT) chloride were from Sigma-Aldrich (St. Louis, MO). All other reagents were from Thermo Fisher Scientific (Suwanee, GA) unless noted.

### Cell culture

Primary bone marrow cultures were prepared from 8-weeks old, C57BL/6J female mice (RRID:IMSR_JAX:000664,Jackson Laboratories, Bar Harbor, ME). Mice were housed 4-5 per cage, with a 12 hour light cycle. Water and food (2018 Teklad Global 18% Protein Rodent Diet, Irradiated, Harlan Laboratories, Indianapolis, IN) were provided *ad libitum*. The mice were euthanized for collection of bone marrow two days after arrival. After euthanasia (cervical dislocation under terminal euthanasia followed by pneumothorax), the limbs were aseptically dissected, and soft tissue was removed from the bone. The bone marrow was flushed from the femur, tibia and humerus, and then strained through a 70 μm cell strainer. The flushed cells were diluted in MSC media (α-MEM containing 10% FBS, 100 U/ml penicillin, 100 μg/ml streptomycin, 0.25 μg/ml amphotericin B), and cells from 2-4 animals were pooled and plated so that each pool represented “n” experiment. Cells were seeded at 6×10^6^/ml in 2 ml per well in 6-well plates. Medium was replaced 5 days after plating, and differentiation was induced at day 7. Prior to the day 7 medium change, undifferentiated cells were harvested as a control for gene expression analysis. For microarray experiments, the medium was replaced with osteoinductive medium consisting of α-MEM, 12.5 μg/ml l-ascorbate, 8 mM β-glycerol phosphate, 0.5 μg/ml insulin, 10 nM dexamethasone, 100 U/ml penicillin, 100 μg/ml streptomycin. Upon addition of differentiation medium, cells received no treatment (naïve) or were dosed with vehicle (DMSO, 0.1% final concentration), rosiglitazone (Rosi, 100 nM), LG100268 (LG268, 100 nM) or TBT (80 nM). Following 4 days of incubation (medium was replaced and cultures were re-dosed once), cells were harvested for gene expression.

For gene and phenotypic validation experiments, BM-MSC and 3T3 L1 (ATCC Cat# CL-173, RRID:CVCL_0123, Lot # 63343749) cells were cultured with adipocyte differentiation medium consisting of DMEM, 10% FBS, 250 nM dexamethasone, 167 nM of 1 μg/ml human insulin, 0.5 mM IBMX, 100 U/ml penicillin, 100 μg/ml streptomycin. Upon addition of differentiation medium, cells received no treatment (naïve) or were dosed as above. On day 3 of differentiation, medium was replaced adipocyte maintenance medium (DMEM, 10% FBS, 250 nM dexamethasone, 167 nM human insulin, 100 U/ml penicillin, 100 μg/ml streptomycin), and the cultures were re-dosed. On days 5 and 7 of differentiation, the adipocyte maintenance medium was replaced and the cultures re-dosed. Following 5 or 10 days of differentiation, cells were harvested for analysis of mRNA expression, lipid accumulation, and mitochondrial biogenesis analyses.

### Microarray data preprocessing and filtering

Microarray data processing and analyses were performed in R software (version 3.3.1). Affymetrix CEL files were processed using the Bioconductor package *oligo*, with the function read.celfiles. Gene expression was normalized using the package *rma*, which performs background correction and quantile normalization. Bioconductor annotation packages *mogene20sttranscriptcluster.db* and *pd.mogene.2.0.st* were used to annotate at the transcript (probeset) level. Data filtering consisted of removal of probesets without mapping to gene symbols, which ultimately led to 24,996 probesets retained for differential expression analysis.

### Assessment of absolute expression of nuclear receptor genes

The absolute expression of nuclear receptors (e.g. PPARγ, RXR and its nuclear receptor binding partners) was estimated in (Day 0) naïve cells, using the method by Piccolo et al. (2013), which estimates the probability of absolute expression of the nuclear receptors at Day 0 (Piccolo et al. 2013). The function UPC() from the SCAN.UPC R package was used to estimate absolute gene quantification and GC content bias was accounted for in the absolute gene expression estimation.

### Differential gene expression analyses

Differential analyses were performed on the normalized expression data using a two-group moderated t-test as implemented in the Bioconductor limma package (Ritchie et al. 2015). Heatmaps depicting significant differentially expressed genes were generated with rows and columns clustered based on Pearson correlation and Euclidean distance, respectively. The Ward agglomeration rule was used for both row and column clustering. Venn diagrams were created using the number of differentially expressed genes with false discovery rate (fdr) < 0.05 for each set of comparisons: each chemical (Rosi, LG268, and TBT) vs. the vehicle (Vh) in differentiated BM-MSCs.

### Pathway enrichment analyses

Pathway enrichment analyses were performed using a hypergeometric distribution-based test to determine the gene sets (Gene Ontology terms from GO database) over-represented in the lists of significant (fdr< 0.05) differentially expressed genes. Top enriched gene sets between each chemical vs. vehicle comparison included pathways related to brown adipocyte differentiation or mitochondrial biogenesis. We curated two separate lists of genes related to brite adipocyte differentiation or mitochondrial biogenesis and related to osteogenesis to validate the upregulation and downregulation of these pathways with our lists of differentially expressed genes and associated t-test statistics (see Table S2).

Validation of the enrichment of these curated gene sets was performed based on the Gene Set Enrichment Analysis (GSEA) software (Subramanian et al. 2005). Further, we validated the observations from our differential gene and pathway analyses in BM-MSCs exposed to Rosi or TBT with publicly available data (GEO Accession: GSE53004) that examined Rosi or TBT exposures in 3T3-L1, another cell model commonly used in adipocyte biology (Pereira-Fernandes et al. 2014).

### mRNA Expression

Total RNA was extracted and genomic DNA was removed using the RNeasy Plus Mini Kit (Qiagen, Valencia, CA). Microarray analyses were performed by the Boston University Microarray and Sequencing Resource using GeneChip^®^ Mouse Gene 2.0ST arrays (Affymetrix, Santa Clara, CA). For RT-qPCR analyses, cDNA was prepared from total RNA using the GoScript™ Reverse Transcription System (Promega), with a 1:1 mixture of random and Oligo (dT)15 primers. All qPCR reactions were performed using the GoTaq^®^ qPCR Master Mix System (Promega). The qPCR reactions were performed using a 7500 Fast RealTime PCR System (Applied Biosystems, Carlsbad, CA): hot-start activation at 95°C for 2 min, 40 cycles of denaturation (95°C for 15 sec) and annealing/extension (55°C for 60 sec). The primer sequences are provided in Table S1. Relative gene expression was determined using the Pfaffl method to account for differential primer efficiencies (Pfaffl 2001), using the geometric mean of the Cq values for beta-2-microglobulin (*B2m*) and 18s ribosomal RNA (*Rn18s*) for normalization. The Cq value from naϊve, undifferentiated cultures was used as the reference point. Data are reported as “Relative Expression”, unless log-transformed and reported as “Log Relative Expression.”

### Phenotype Analyses

Analyses were carried out following 10 days of differentiation and treatment of 3T3 L1 cells. To determine lipid accumulation, the cells were washed with PBS and incubated with Nile Red (1 μg/ml in PBS) for 15 minutes, in the dark, at room temperature. Fluorescence (excitation 485 nm, 20 nm bandwidth; emission 530 nm, 25 nm bandwidth) was measured using a BioTek Synergy2 plate reader (Biotek Inc., Winooski, VT). The fluorescence in experimental wells was normalized by subtracting the fluorescence measured in naïve (undifferentiated) cells and reported as “RFUs.” To measure mitochondrial biogenesis, the MitoBiogenesis In-Cell Elisa Colorimetric Kit (Abcam, Cambridge, UK, #110217) was used, following the manufacturer’s standard protocol. The levels of two mitochondrial proteins (COX1 and SDH) that are measured simultaneously in each well and normalized to the total protein content via JANUS staining. Absorbance (OD 600nm for COX1, OD 405nm for SDH, and OD 595nm for JANUS) was measured using a BioTek Synergy2 plate reader. The absorbance ratios of COX/SDH in experimental wells were normalized to the naïve (undifferentiated) cells.

### Statistics

All statistical analyses were performed with Prism 5 (GraphPad Software, Inc., La Jolla, CA). Data are presented as means ± standard error (SE). Four biological replicates of naïve, vehicle, or each chemical-treated cells were processed for the microarray. The n value indicates the number of independent samples that were evaluated. The qPCR data were log-transformed before statistical analyses. One-factor ANOVAs (Dunnett’s) were performed to analyze the qPCR and phenotypic data.

## Results

### Nuclear receptor expression in undifferentiated BM-MSCs

BM-MSCs are multipotent cells that can differentiate into adipocytes, chondrocytes, and osteocytes (Pittenger et al. 1999). We began by determining the probability of absolute expression of nuclear receptors in undifferentiated BM-MSCs. Bone marrow cells isolated from female C57BL/6J mice were cultured in basal medium for 7 days. RNA was isolated and analyzed for mRNA expression by microarray. Using the absolute gene quantification in microarrays method by Piccolo et al. (2013), we determined that 8 RXR-related nuclear receptors had greater than a 50% probability of expression in undifferentiated cells: *Lxrb*, *Nur77*, *Pparg*, *Ppard*, *Rara*, *Rxra*, *Rxrb*, and *Vdr* (Figure 1). Thus, at the onset of exposure, TBT activate could PPARγ and RXRs directly (Grün et al. 2006; le Maire et al. 2009), as well as LXRβ, Nur77 and PPARδ through permissive activation via RXR (Giner et al. 2015).

**Figure 1.**
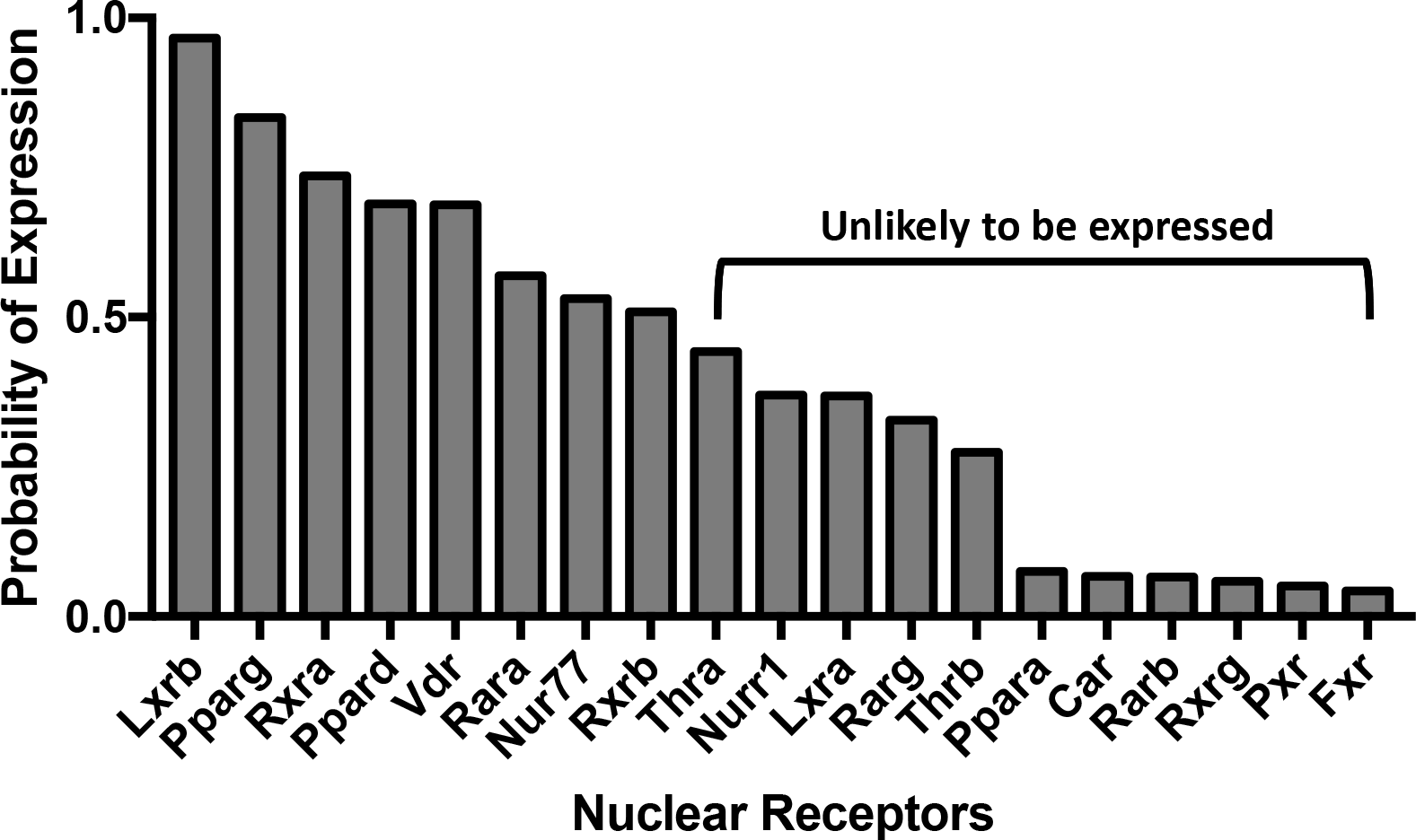
Expression of retinoid X receptors (RXR) and their nuclear receptor partners in BM-MSCs. RNA was isolated from bone marrow cells isolated from female, 8 week old, C57BL/6J mice and cultured in basal medium for 7 days. mRNA expression was analyzed by microarray. The probability of expression was determined using the method of Piccolo et al. (2013).

### Chemical treatment modifies nuclear receptor and coregulator expression

Our previous work showed that TBT is able not only to stimulate adipocyte differentiation in BM-MSCs, but also to divert osteogenic differentiation toward adipocyte differentiation (Baker et al. 2015). To determine the potential nuclear receptor pathways involved, BM-MSC cultures were stimulated to undergo osteogenic differentiation and treated with Vh or TBT (80 nM). This concentration falls between the ED_50_ and the maximally efficacious concentration determined in multiple human and mouse cell culture models (Biemann et al. 2012; Carfi et al. 2008; Hiromori et al. 2009; Kirchner et al. 2010; Pereira-Fernandes et al. 2013; Yanik et al. 2011), and is a maximally efficacious concentration in mouse-derived BM-MSCs (Yanik et al. 2011). Furthermore, this concentration of TBT is in line with butyltin concentrations measured in human serum (Kannan et al. 1999). We compared the TBT-induced transcriptional response to those induced by agonists specific for either PPARγ (rosiglitazone (Rosi), 100 nM) or RXR (LG100268 (LG268), 100 nM) at the lowest concentrations shown to be maximally efficacious (Lala et al. 1996; Lehmann et al. 1995). After 4 days of differentiation, RNA was isolated and analyzed for mRNA expression by microarray.

Chemical treatment modified the expression of nuclear receptors that heterodimerize with RXR. In BM-MSCs exposed to Rosi or TBT, expression of nuclear receptors *Pparg*, *Thrb*, *Rarb*, *Rxrg*, *Ppara*, and *Rxra* was greater than in undifferentiated and Vh-treated BM-MSCs (Figure 2a). Rosi and TBT also decreased the expression of *Nr4a1* (*Nur77*) and *Vdr* (Figure 2a). In LG268-treated cells, *Thrb, Rarb*, and *Rarg* were significantly induced and *Nr1i3* (*Car*) was differentially downregulated, compared to undifferentiated BM-MSCs (Figure 2a). To examine a broader spectrum of nuclear receptors, including steroid receptors (Robinson-Rechavi 2003), we queried the data for 47 mouse nuclear receptors identified from the Nuclear Receptor Signaling Atlas (NURSA) (Becnel et al. 2015). Overall, TBT induced a nuclear receptor expression profile similar to both Rosi (PPARγ) and LG268 (RXR) (Figure S1).

**Figure 2.**
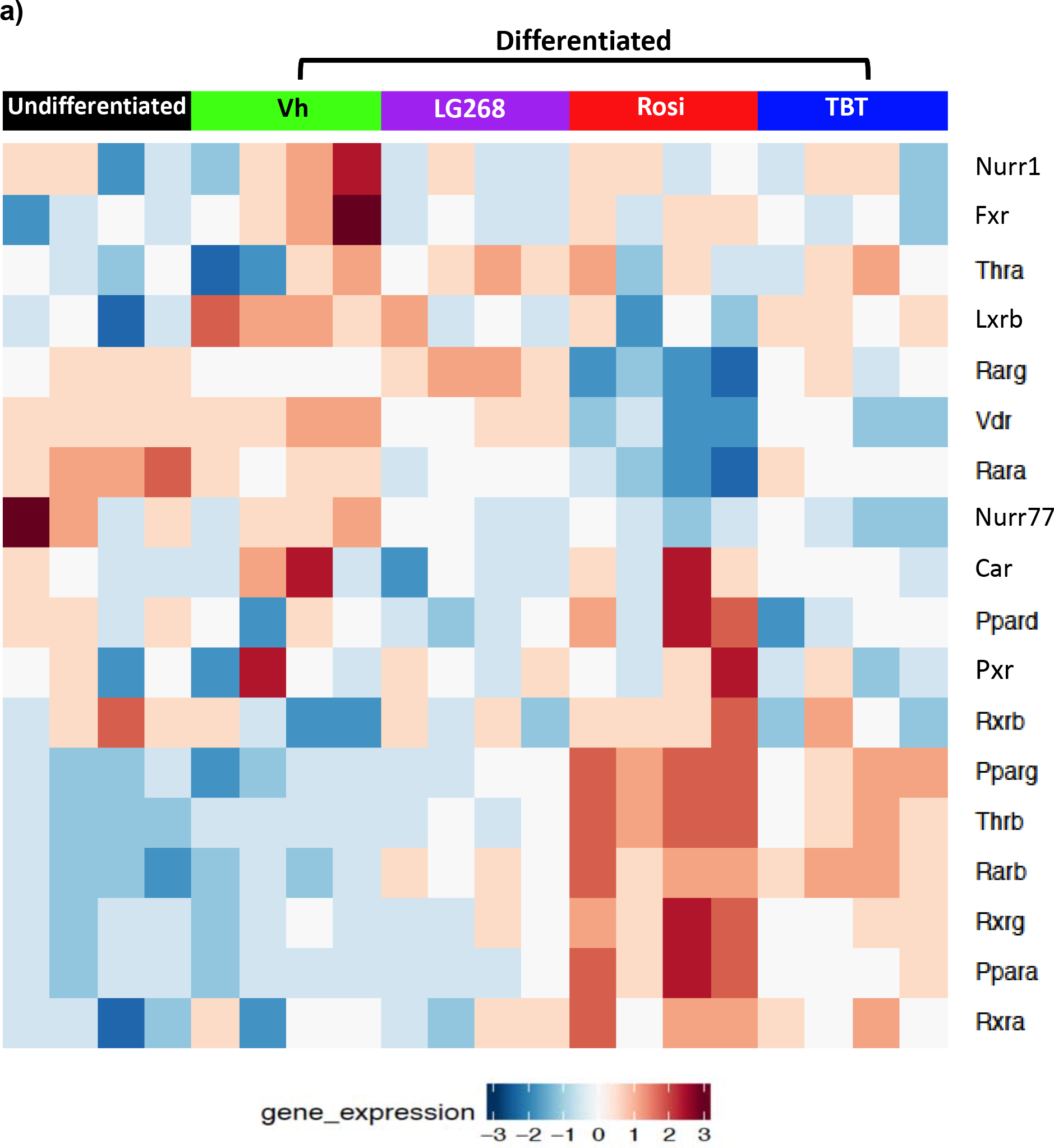
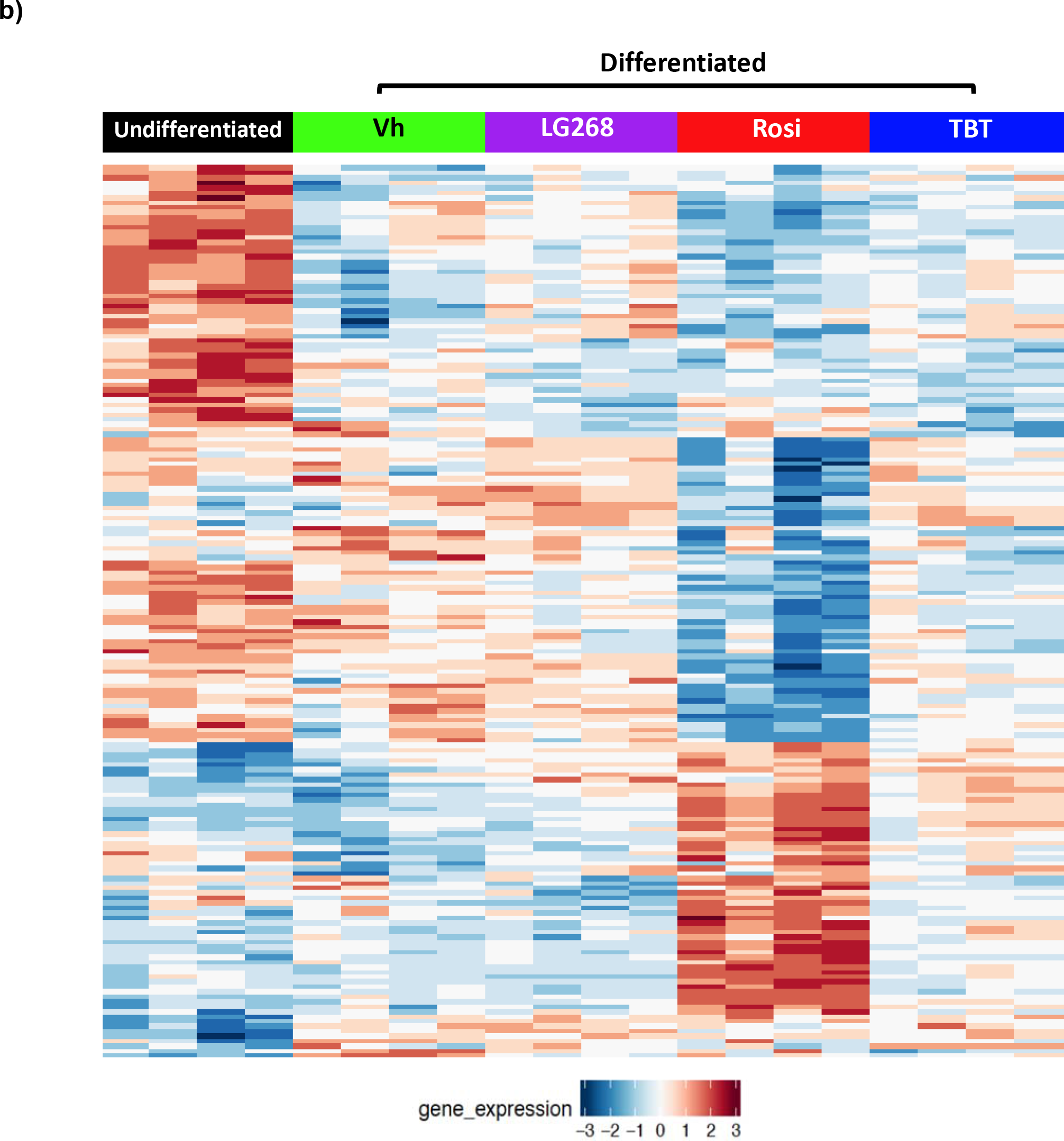
Differential expression of RXR heterodimer partners (A) and nuclear receptor coregulators (B) in BM-MSCs undergoing osteogenic differentiation and treated with PPAR/RXR ligands. Primary bone marrow cells were isolated from female, 8 week old, C57BL/6J mice, plated, and allowed to adhere for 7 days, and undifferentiated cultures were harvested at this time. In experimental cultures, the medium was replaced with basal medium supplemented with osteogenic additives, β-glycerol phosphate, ascorbate, insulin and dexamethasone. Cultures were treated with Vh (DMSO, 0.1%), Rosiglitazone (Rosi, 100 nM), LG100268 (LG268, 100 nM) or TBT (80 nM). After 4 days of culture, cells were harvested and analyzed for gene expression using microarray. The heatmaps display the significant differentially expressed **(A)** nuclear receptors and **(B)** coregulators (fdr<0.05).

Nuclear receptor coregulators play an important role in regulating nuclear receptor transcriptional activities (Feige and Auwerx 2007). Therefore, we examined the expression of nuclear receptor coregulators in the cultures. Using a list of 280 identified mouse nuclear receptor coregulators from NURSA, we determined that there were 181 nuclear receptor coregulators that were significantly differentially expressed between the undifferentiated and chemically-treated BM-MSCs (Figure 2b). TBT has similar coregulator expression profile with Rosi; yet, Rosi induced more strongly some of the coregulator genes’ expressions as indicated by the color intensity of the induced expression (Figure 2b). Thus, we examined more closely the expression profiles of nuclear receptor coregulators by directly comparing Rosi and TBT (Figure S2). The expression pattern of the nuclear receptor coregulators differed significantly (fdr< 0.05) between the two chemicals. Rosi significantly upregulated the expression of coregulators involved in brite/brown adipocyte differentiation (*Ppargc1a*, also known as *Pgc1a*) and mitochondrial function (e.g. *Phb2*). Thus, TBT did not appear to efficiently induce the expression of coregulators involved in brite adipocyte differentiation compared to a therapeutic chemical, rosiglitazone.

### TBT induces a distinct transcriptional response

Comparisons of the overall differentially expressed genes between vehicle (Vh) and chemical (Rosi, TBT, LG268) treatments in differentiated BM-MSCs reveal both overlapping and unique genes across all treatments (Figure 3a). Principal component analyses (PCA) revealed that the Rosi-induced gene expression patterns cluster separately from those induced by TBT and LG268 (Figure 3b). TBT has gene expression profiles in common with both the PPARγ ligand (Rosi) and the RXR ligand (LG268). Using fdr<0.05, there were a total of 4318 genes uniquely up- and down-regulated by Rosi, 274 genes uniquely regulated by TBT, and 18 genes uniquely regulated byLG268 (Figure 3c). There were a total of 164 significantly induced genes in common between all three chemicals. As expected, a set of genes commonly downregulated by all three chemicals were those related to osteogenesis (Figure S3). However, the enrichment of this set had much higher significance for Rosi and TBT (q<0.001) than for LG268 (q=0.05).

**Figure 3.**
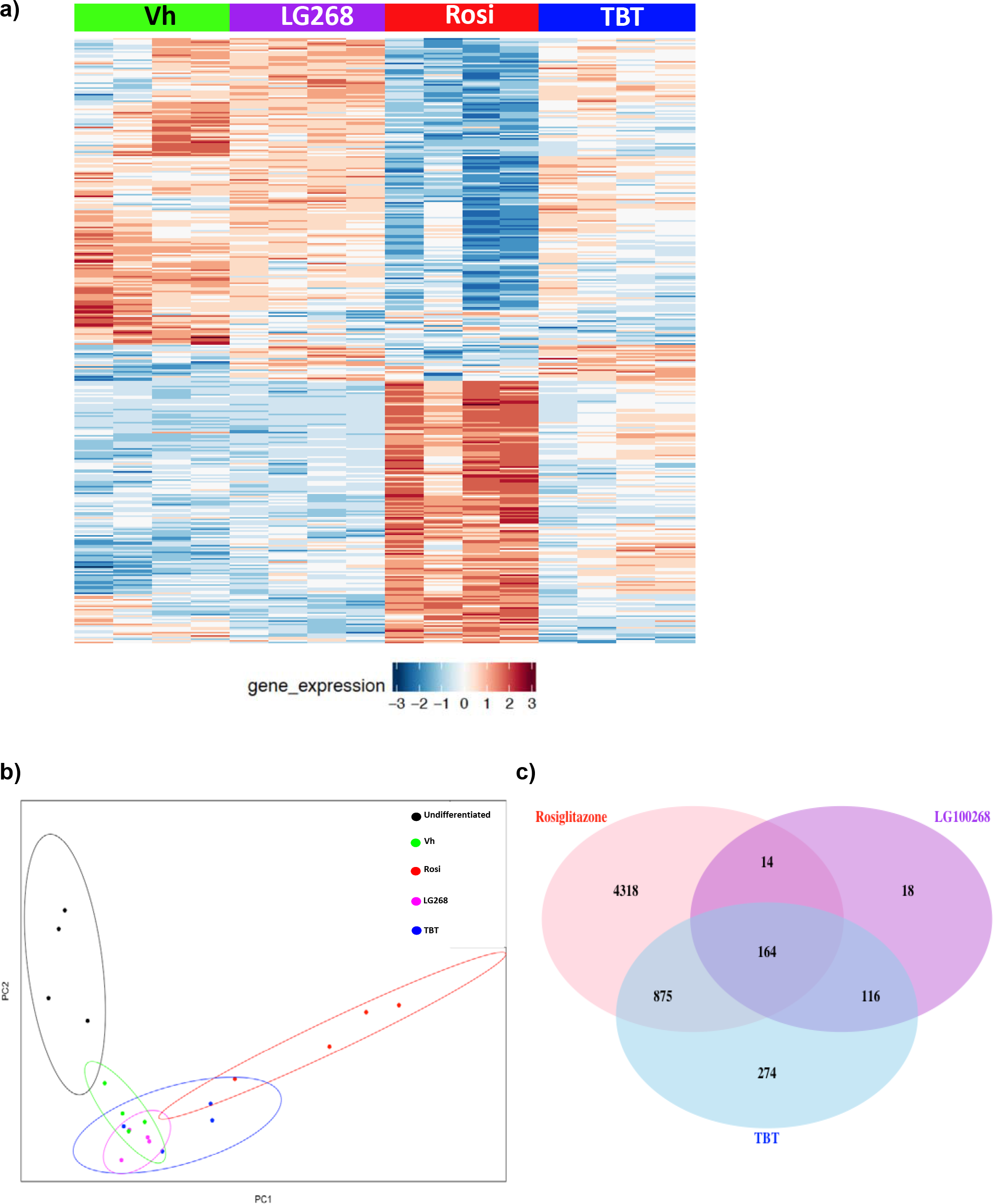
Significant differentially expressed genes induced by rosiglitazone, LG100268 and TBT. Transcriptomic profiles from each chemical (TBT, Rosi, and LG268) were compared to vehicle (DMSO).**(A)** The heatmap depicts 7044 significant differentially expressed genes with false discovery rate (fdr) < 0.05 between the chemicals and Vh, with rows and columns sorted according to hierarchical clustering (as described in the Methods). **(B)** Principal component analyses (PCA) were performed in R, and the plots show the clustering of the undifferentiated and chemically treated cells by the first two principal components. **(C)** The Venn diagram depicts the number of significant differentially expressed genes with fdr<0.05 for each set of chemical comparisons.

The significant differential genes for each chemical (compared to Vh-treated cells) were used for pathway enrichment analyses using GO (Biological Processes) terms. The most significantly enriched biological processes in Rosi-treated cells were pathways related to brown adipocyte differentiation or mitochondrial biogenesis (Figure 4). The most significantly enriched pathways in TBT-treated BM-MSCs were pathways related to lipid metabolic processes as well as to metabolic processes (Figure 4). Biological processes such as cell cycle regulation were significantly enriched in cells treated with LG268 (Figure 4). The overall pathways common and different between all chemicals are shown in Figure S4. Ultimately, the environmental chemical, TBT, does not induce a gene expression pattern specifically like the PPARγ ligand or the RXR ligand.

**Figure 4.**
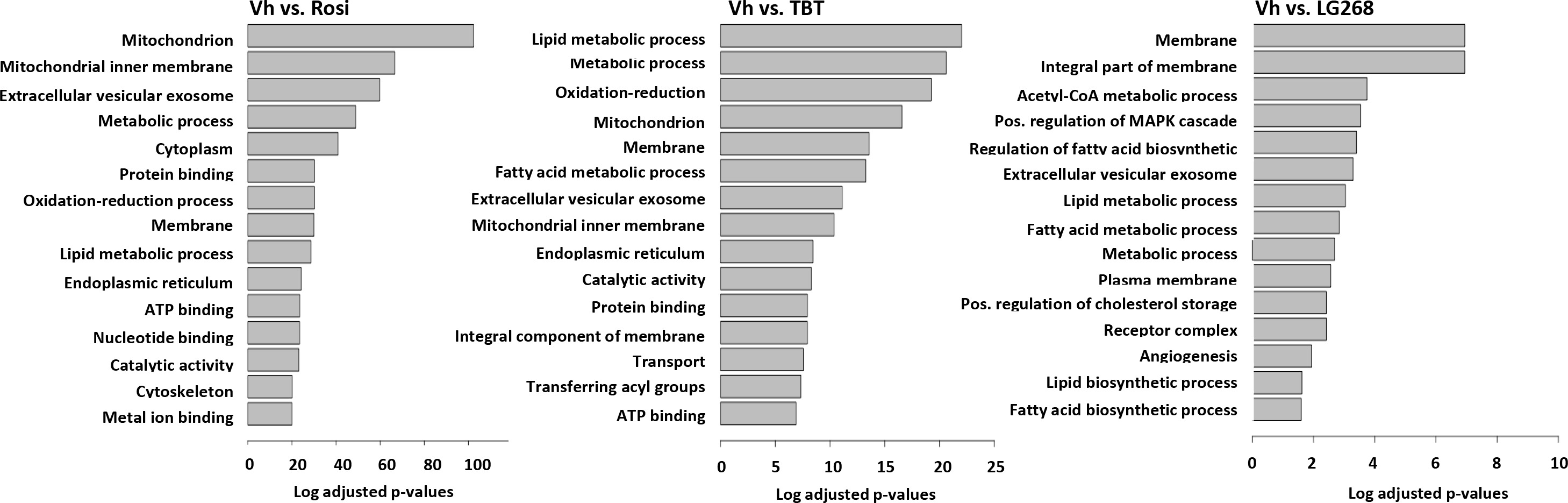
Top enriched pathways (GO Terms) induced by rosiglitazone (A), LG100268 (B) and TBT (C). Pathway enrichment analyses were performed using a hypergeometric distribution based test to determine the gene sets (Gene Ontology terms from GO database) over-represented in the lists of significant (fdr< 0.05) differentially expressed genes.

### TBT does not efficaciously induce health-promoting genes

We directly compared the enriched pathways between Rosi and TBT and found that mitochondrial biogenesis was significantly upregulated by Rosi compared to TBT (Figure 5a). To investigate the difference between the transcriptional responses more thoroughly, we curated gene sets, from the literature, related to brite/brown adipocyte differentiation and mitochondrial biogenesis (Table S2). Using GSEA software, we determined which curated pathways were enriched in Rosi- versus TBT-treated cultures. Rosi effectively upregulated metabolism- and mitochondria-related pathways in differentiated BM-MSCs, with enrichment score (ES) of 0.82 (q<0.001) and 0.80 (q<0.001), respectively (Figure 5b; Figure S5a). Other significant pathways induced by Rosi were transcriptional factors/coregulators and brite/brown adipocyte differentiation (Figure 5b). To validate whether the genes in the curated genesets related to brite adipocyte differentiation and mitochondrial biogenesis were uniquely upregulated by Rosi, we compared the differential expression of these genes across the three chemicals. Rosi significantly induced most of the genes (e.g. *Coq3*, *Coq9*, *Acaa2*, *Pck1*) related to mitochondria and metabolism as highlighted in boxed area “a”, and TBT-treated cells induced some genes in common with LG268 that were immune- or inflammatory-related (e.g. *Ifnar2*, *IL10*, *Kng1*) and hormone/vitamin (e.g. *Irs1, Cnp*) as shown in boxed area “b” (Figure 5c).

**Figure 5.**
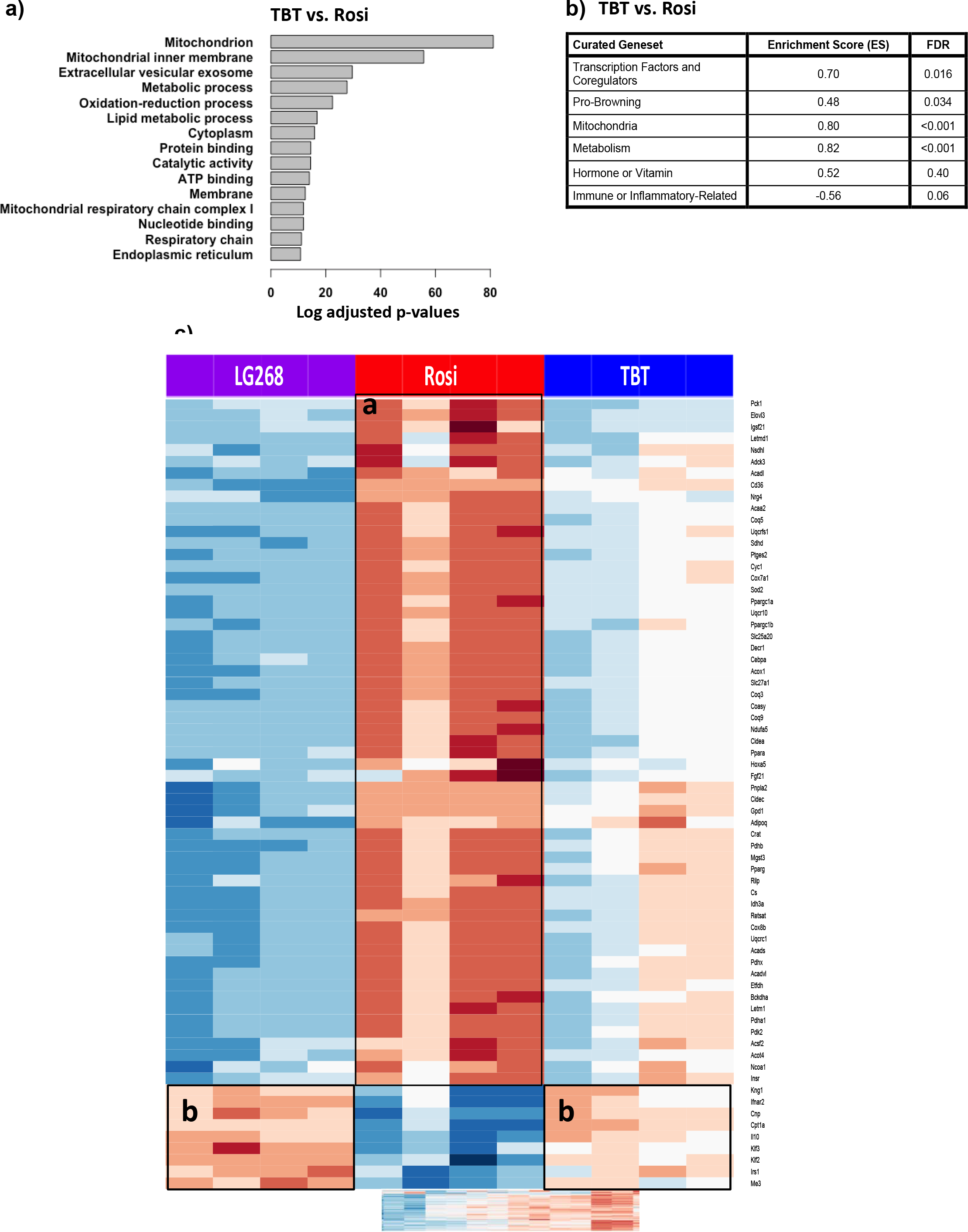
Pathway enrichment analyses related to brite/brown adipogenesis and mitochondrial biogenesis. **(A)** Most significant differentially enriched pathways between the Rosi- and TBT-treated cells. **(B)** GSEA analysis of curated genesets related to brown adipocyte differentiation and mitochondrial biogenesis.**(C)** The heatmap displays the significant differentially expressed genes (fdr<0.05) related to mitochondrial biogenesis and adipocyte browning between Rosi, LG200268 and TBT.

We used two approaches to validate the finding that TBT induces a transcriptional response distinct from Rosi. First, BM-MSCs were cultured for 5 and 10 days in adipogenic medium to evaluate effects on adipocyte differentiation by the three chemicals. We analyzed by RT-qPCR expression of genes common to all classes of adipocytes (Figure 6a), as well as those involved in brite/brown adipogenesis (Figure 6b). Rosi and TBT both induced *Pparg* expression, with Rosi more robustly inducing expression than TBT (Figure 6a). The expression levels of adipogenic target genes of PPARγ, such as *Fabp4* (Fatty Acid Binding Protein 4), *Plin1* (Perilipin 1), and *Scd1* (Stearoyl-Coa Desaturase 1), were all significantly induced by TBT and Rosi by day 10 of differentiation (Figure 6a). Compared to TBT and LG268, Rosi more strongly induced the nuclear receptor, *Ppara*, and coregulator, *Pgc1a* (peroxisome proliferator-activated receptor gamma coactivator 1-alpha), which play roles in regulating mitochondrial biogenesis and brite/brown adipocyte differentiation, respectively (Figure 6b). Additionally, brite adipocyte-related genes *Cidea* (Cell Death-Inducing DFFA-like Effector A) and *Elovl3* (Elongation of Very Long Chain Fatty Acid 3) were uniquely and significantly induced by Rosi (Figure 6b). The classic marker of brite/brown adipocytes, *Ucp1* (Uncoupling protein 1), showed a trend toward upregulation by Rosi, but the expression level did not reach statistical significance because of high variability (Figure 6b).

**Figure 6.**
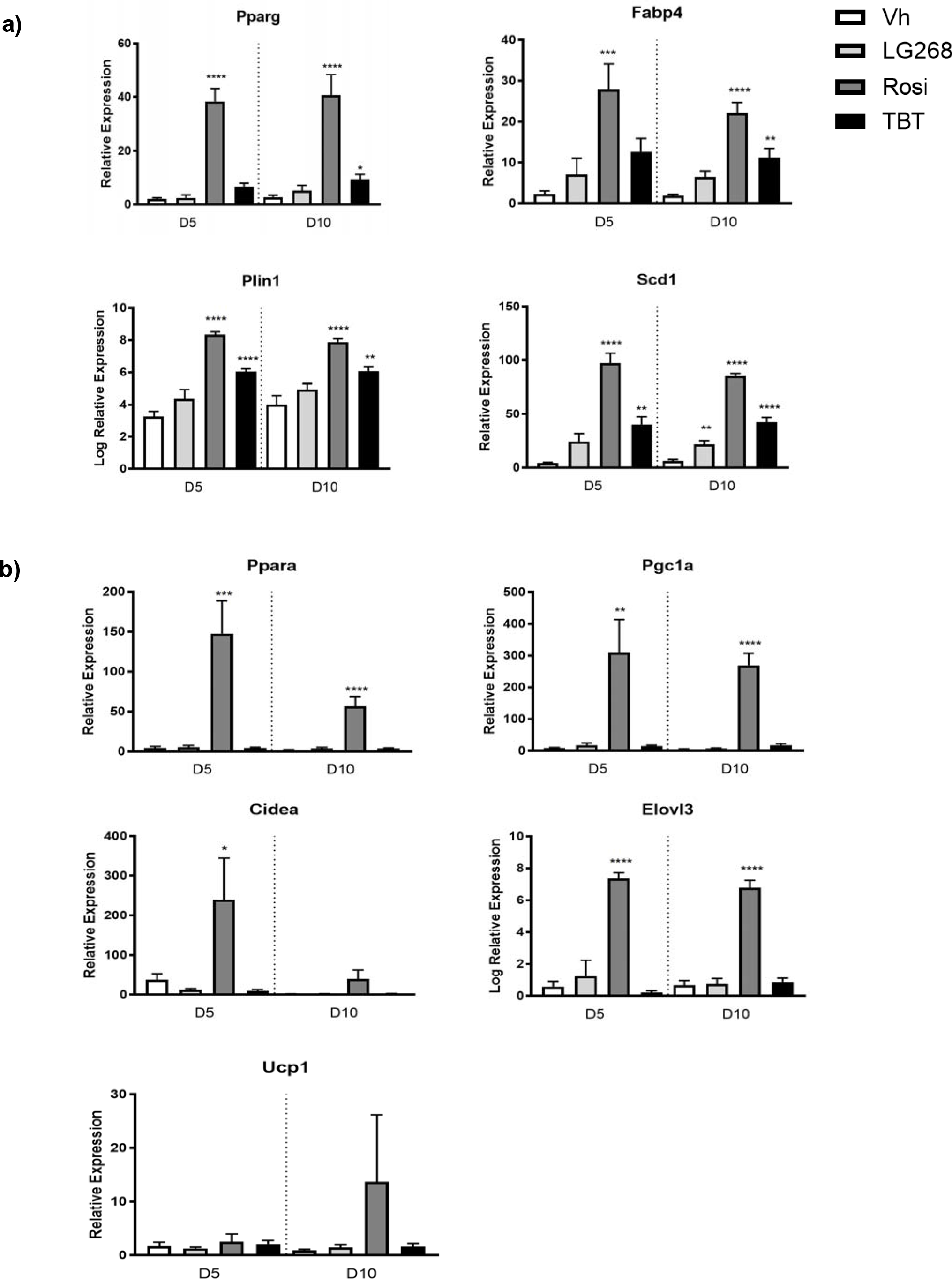
Differential adipogenic gene expression induced by rosiglitazone versus TBT in BM-MSCs. Primary bone marrow cells were isolated from female, 8 week old, C57BL/6J mice, plated, and allowed to adhere for 7 days. The medium was replaced with DMEM supplemented with adipogenic additives: 10% FBS, 250 nM dexamethasone, 167 nM of 1 μg/ml human insulin, 0.5 mM IBMX. Cultures were treated with Vh (DMSO, 0.1%), Rosiglitazone (Rosi, 100 nM), LG100268 (LG268, 100 nM) or TBT (80 nM). On days 3, 5 and 7 days of differentiation, medium was replaced with adipocyte maintenance medium (DMEM, 10% FBS, 250 nM dexamethasone, 167 nM of 1 μg/ml human insulin), and the cultures were re-dosed. Following 5 or 10 days of differentiation, cells were harvested and analyzed for gene expression by RT-qPCR. **(A)** Common PPARγ-related genes. **(B)** Genes related to brite/brown adipogenesis and mitochondrial biogenesis. Statistically different from Vh-treated on the same day (*p<0.05, **p<0.01, ***p<0.001; ****p<0.0001, ANOVA, Dunnett’s).

Second, we validated the unique upregulation of brite adipogenesis and mitochondrial biogenesis by Rosi (and not by TBT) with a publicly available transcriptomic dataset (GSE53004) generated in 3T3 L1 cells and with phenotypic analyses of 3T3 L1 cells, a preadipocytes model commonly used in adipocyte biology (Pereira-Fernandes et al. 2014). Consistent with our observations in Rosi- and TBT-treated BM-MSCs, Rosi upregulated the mitochondrial and metabolism pathways in 3T3 L1 cells (both ES=0.68) more so than TBT (ES=0.87) (Figure 7a, Figure S5). Rosi also significantly induced more genes in the curated pathways related to adipocyte browning and mitochondria than TBT (Figure 7b). Both Rosi and TBT significantly induced lipid accumulation in differentiating 3T3 L1 cells (Figure 8a). However, only Rosi significantly upregulated mitochondrial biogenesis in the differentiating 3T3 L1 (Figure 8b). Thus, the distinct upregulation of adipocyte browning and mitochondrial biogenesis by Rosi is not unique to BM-MSCs.

**Figure 7.**
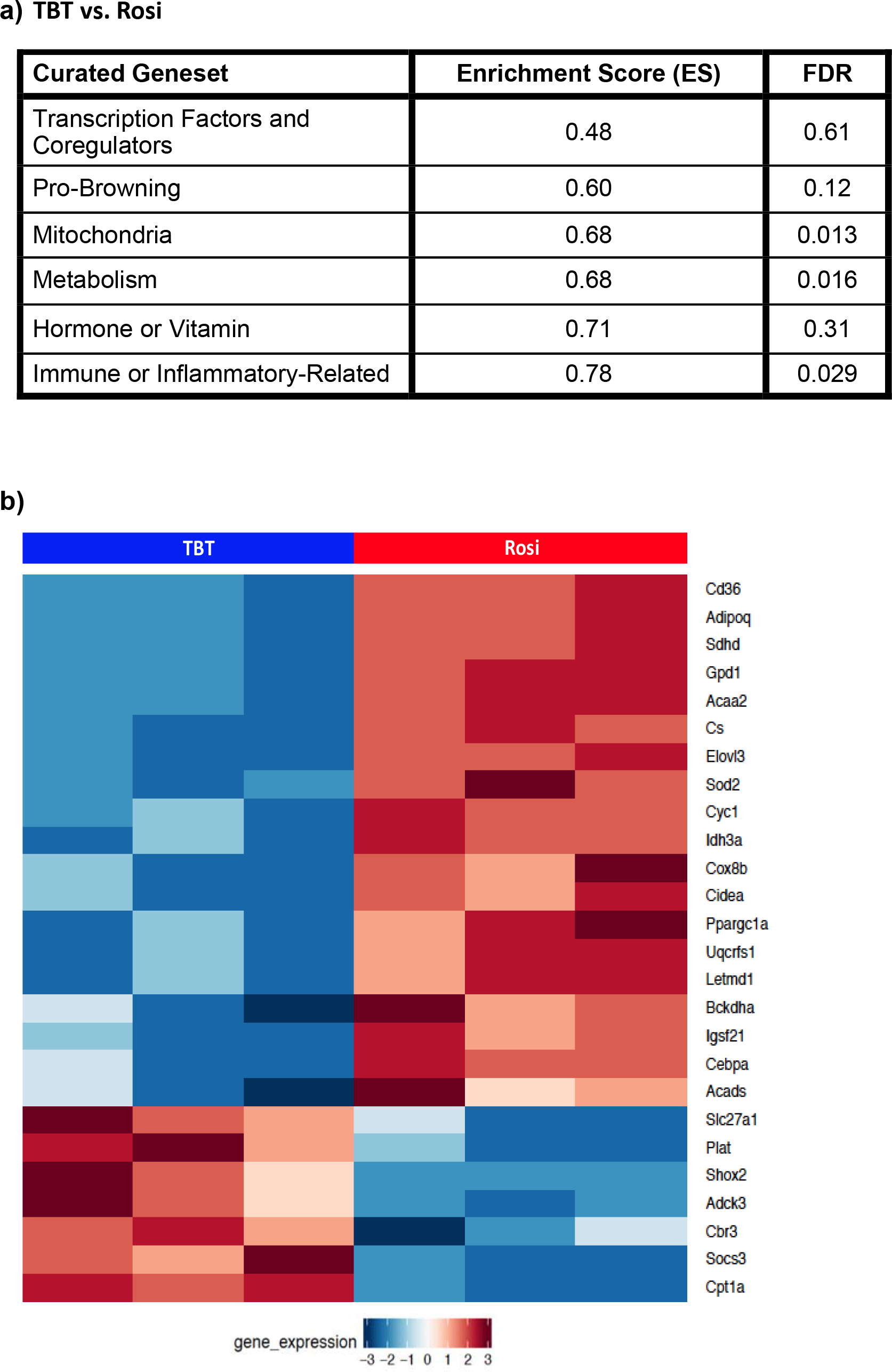
Differential adipogenic gene expression induced by rosiglitazone versus TBT in 3T3 L1 cells. The publicly available dataset (GSE53004) was generated from 3T3-L1 cells treated with Rosiglitazone (500 nM) or TBT (50 nM) for 10 days in DMEM with 10% FBS (Pereira-Fernandes et al. 2014). **(A)** GSEA analysis of curated genesets related to brown adipocyte differentiation and mitochondrial biogenesis. **(B)** The heatmap displays the significant differentially expressed genes (fdr<0.05) related to mitochondrial biogenesis and adipocyte browning between Rosi and TBT.

**Figure 8.**
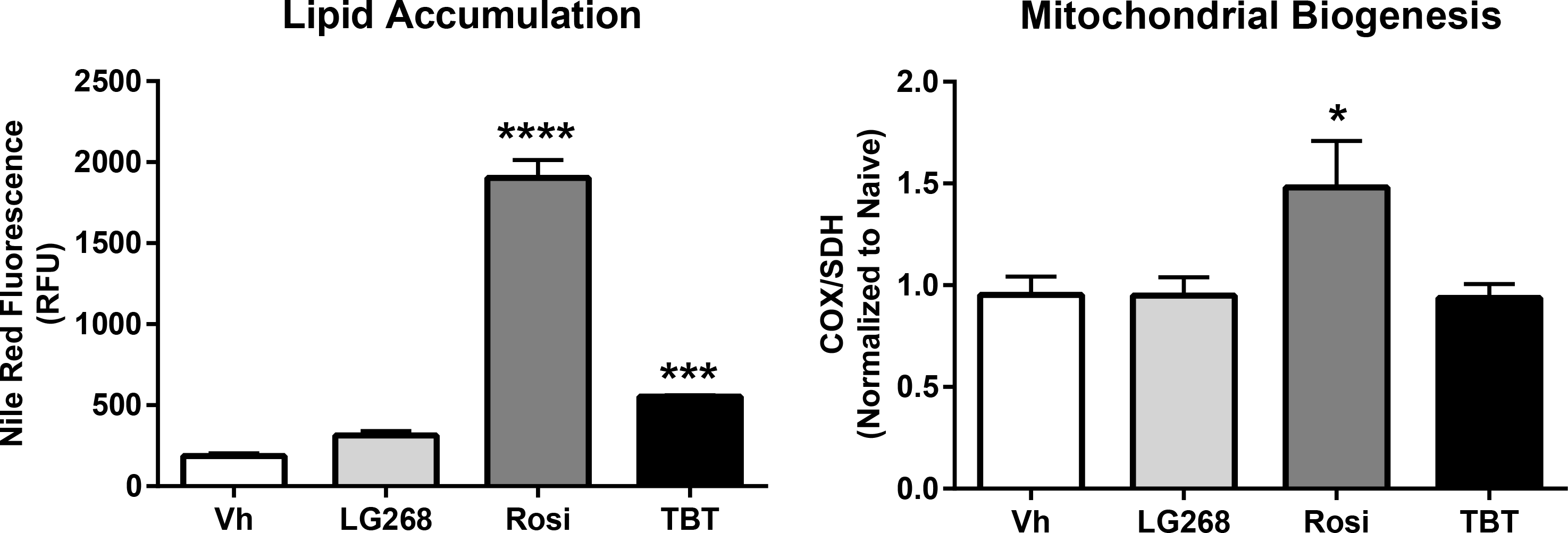
Lipid accumulation and mitochondrial biogenesis induced by rosiglitazone versus TBT in 3T3 L1 cells. Cells were plated and then incubated for 4 days. Differentiation and dosing were carried out as described in Figure 6. Following 10 days of differentiation, cells were washed and analyzed for either **(A)** lipid accumulation (Nile Red staining) or **(B)** mitochondrial biogenesis (ELISA). Statistically different from Vh-treated on the same day (*p<0.05, ***p<0.001; ****p<0.0001, ANOVA, Dunnett’s).

## Discussion

TBT, the original “environmental obesogen,” binds and activates both PPARγ and RXRs, and its engagement of several nuclear receptor pathways contributes to the effects of this environmental toxicant on adipose and bone homeostasis (Baker et al. 2015; Grün et al. 2006; le Maire et al. 2009; Watt and Schlezinger 2015; Yanik et al. 2011). We used primary BM-MSCs to investigate the molecular pathways that are activated and/or disrupted by TBT during osteoblast and adipocyte differentiation. Because of TBT’s ability to interact directly with both PPARγ and RXR, we included specific ligands for these receptors: rosiglitazone (PPARγ ligand) and LG100268 (RXR ligand) for comparison. Here, we show that, as expected, rosiglitazone, TBT and LG100268 all suppress osteogenesis; but that unexpectedly, TBT has a limited capacity to fully activate PPARγ’s transcriptional programs that regulate energy homeostasis.

There has been a challenge in understanding the potential for obesogenic effects of environmental PPARγ ligands (e.g. TBT) to lead to adverse effects on metabolic health, as people treated with therapeutic PPARγ ligands (e.g. rosiglitazone) gain weight but improve their metabolic health (Soccio et al. 2014). Recent studies support the idea that, “PPARγ agonism is not correlated directly with anti-diabetic action” (Choi et al. 2014). The biological pathways regulated by PPARγ (e.g. white adipogenesis, brite/brown adipogenesis, insulin sensitivity suppression of osteogenesis) are regulated separately through differential coregulator recruitment and post-translational modifications (Choi et al. 2010, 2011; Claussnitzer et al. 2015; Cohen et al. 2014; Cohen and Spiegelman 2015; Feige and Auwerx 2007; Timmons and Pedersen 2009; Wang et al. 2016). The unliganded PPARγ/RXR receptors are engaged in large complexes of corepressors and coactivators that can promote repression or activation of PPARγ target genes (Guan 2005; Michalik et al. 2007; Tudor et al. 2007). In order to induce PPARγ transcriptional activation, ligand binding initiates corepressor release and coactivator recruitment through protein structure remodeling and epigenetic changes of the chromatin (Feige and Auwerx 2007). Furthermore, acetylation and dephosphorylation of PPARγ are crucial to the release and recruitment of its coregulators. For example, phosphorylation of S273 of PPARγ by CDK5 or ERK not only induces a “diabetogenic” program of gene expression by attenuating expression of insulin sensitizing genes but also reduces brite/brown gene expression (Banks et al. 2015).

Rosiglitazone induced a distinct spectrum of transcriptional coregulators, relative to TBT. In particular, TBT does not efficaciously induce expression of coregulators related to mitochondrial biogenesis such as *Phb2* (see Figure S2). Phb2 is required for modulation of mitochondrial assembly and respiration (Pbm et al. 2016). Rosiglitazone also uniquely induced expression of *Pgc1a*, which is a crucial transcriptional coregulator of brite/brown fat differentiation (Feige and Auwerx 2007; Koppen and Kalkhoven 2010; Seale 2015). PGC1α is also an important transcriptional inducer of mitochondrial biogenesis as it plays a role in regulating transcription of nucleus encoded mitochondrial genes and activates other transcription factors (e.g. NRF-1 and NRF-2) that influence expression of key mitochondrial enzymes such as ATP synthetase and cytochrome *c* oxidase (Aquilano et al. 2010; Jornayvaz and Shulman 2010). Another environmental PPARγ ligand, MEHP, modifies PPARγ coregulator recruitment distinctly from rosiglitazone (Feige et al. 2007). While MEHP binds to PPARγ in a configuration similar to rosiglitazone, MEHP is unable to stabilize helix 12 of PPARγ thus limiting its ability to release the corepressor NCOR, and MEHP stimulates the recruitment of the coactivators MED1 and PGC1α but not of p300 and SRC1 (Feige et al. 2007). Therefore, TBT may not only differentially induce coregulator expression, but we also hypothesize that TBT, like MEHP, is a selective PPARγ ligand, with a limited capacity to release and recruit coregulators.

Moreover, TZDs such as pioglitazone and rosiglitazone can induce mitochondrial biogenesis and fatty acid oxidation in human adipose tissue (Harms and Seale 2013). We observed upregulation of genes related to mitochondrial biogenesis (*Ppara* and *Cidea*); browning of fat (*Elovl3* and *Pgc1a*) only by rosiglitazone both in the osteogenic BM-MSC cultures as well as in the adipogenic BM-MSC cultures. Importantly, only rosiglitazone upregulated mitochondrial protein expression in 3T3 L1 cells, as well.

TBT’s limited ability to activate the PPARγ-driven transcriptional repertoire does not only occur in BM-MSCs, but also in the classic adipogenic model, 3T3 L1 cells. Our analysis of the publicly available transcriptional data set derived from 3T3-L1 cells showed that rosiglitazone, but not TBT, was able to efficaciously upregulate expression of genes and pathways related to brite fat differentiation and mitochondrial function. TBT did not induce mitochondrial biogenesis in our study, which is in line with a previous study showing that TBT induces a phenotypically distinct adipocyte in 3T3 L1 cells (Regnier et al. 2015). Furthermore, unlike rosiglitazone, TBT not only induces weight gain *in vivo* but also metabolic disruption (i.e. hepatic steatosis, hyperinsulinemia and hyperleptinemia) following adult exposure (Bertuloso et al. 2015; Oakes et al. 1994; Zuo et al. 2011).

Here, we provide support for the novel hypothesis that environmental PPARγ ligands, such as but not limited to TBT, act as selective PPARγ ligands that favor white over brite adipogenesis. While these studies investigated this hypothesis using *in vitro* mouse models, they are an important step in understanding how white adipocyte- (vs. brite adipocyte-) skewed differentiation may impact human health. First, TBT is well known to induce adipogenesis and metabolic disruption *in vivo* (Bertuloso et al. 2015; Grun et al. 2006; Zuo et al. 2011). Second, there is a high degree of structural and functional similarity between the mouse and human PPARγ’s (Lambe and Tugwood 1996). Last, TBT has been shown to be highly potent and efficacious as stimulating adipocyte differentiation in human adipose- and bone marrow-derived MSCs (Carfi et al. 2008; Kirchner et al. 2010). Therefore, it is highly likely that TBT also is able to skew adipocyte differentiation in human cells, and as human pre-adipocytes become more readily available, we will test our hypothesis in a human system.

Critically, when homeostasis is out of balance, white adipocytes become dysfunctional, this results in adipose stress that leads to lipid overflow, systemic inflammation and insulin resistance (Stinkens et al. 2015; Wensveen et al. 2015). Humans with minimal brite adipocyte populations are at higher risk for obesity and type 2 diabetes (Claussnitzer et al. 2015; Sidossis and Kajimura 2015; Timmons and Pedersen 2009).Thus, it is essential to understand how environmental PPARγ ligands modify not only adipocyte differentiation but also adipocyte function (energy storage vs. insulin sensitization and energy dissipation) in order to assess the risk to metabolic health.

## Conclusions

PPARγ can regulate white and brite/brown adipogenesis independently, with distinct transcriptional profiles controlled through specific ligand binding. Compared to a therapeutic PPARγ ligand, rosiglitazone, the environmental metabolic disruptor, TBT, is a PPARγ ligand that selectively activates PPARγ transcriptional activities that favor white adipogenesis at the expense of promoting the browning and healthy metabolic activities of PPARγ.

## Ethics Statement

Animal studies were reviewed and approved by the Institutional Animal Care and Use Committee at Boston University and performed in an American Association for the Accreditation of Laboratory Animal Care accredited facility (Animal Welfare Assurance Number: A3316-01). All animals were treated humanely and with regard for alleviation of suffering.

## Acknowledgements

The authors would like to thank Ms. Cassie Huang for her superb technical assistance. Analytical assistance was provided by the Bioinformatics and Molecular Modeling Core of the Boston University Superfund Program. This work was supported by a Superfund Research Program grant P42ES007381 (J.J.S, S.M.), and a “Seed the Scientist” award from the Find the Cause Breast Cancer Foundation (A.L, S.M).

## References

Anghel SI, Bedu E, Vivier CD, Descombes P, Desvergne B, Wahli W. 2007. Adipose tissue integrity as a prerequisite for systemic energy balance: a critical role for peroxisome proliferator-activated receptor gamma. J Biol Chem 282:29946–29957; doi:10.1074/jbc.M702490200.

Aquilano K, Vigilanza P, Baldelli S, Pagliei B, Rotilio G, Ciriolo MR. 2010. Peroxisome proliferator-activated receptor gamma co-activator 1alpha (PGC-1alpha) and sirtuin 1 (SIRT1) reside in mitochondria: possible direct function in mitochondrial biogenesis. J Biol Chem 285:21590–21599; doi:10.1074/jbc.M109.070169.

Baker AH, Watt J, Huang CK, Gerstenfeld LC, Schlezinger JJ. 2015. Tributyltin engages multiple nuclear receptor pathways and suppresses osteogenesis in bone marrow multipotent stromal cells. Chem Res Toxicol 28:1156–1166; doi: 10.1021/tx500433r.

Baker AH, Wu TH, Bolt AM, Gerstenfeld LC, Mann KK, Schlezinger JJ. 2017. Tributyltin alters the bone marrow microenvironment and suppresses B cell development. Toxicol Sci Off J Soc Toxicol; doi: 10.1093/toxsci/kfx067.

Banks AS, McAllister FE, Camporez JPG, Zushin P-JH, Jurczak MJ, Laznik-Bogoslavski D, et al. 2015. An ERK/Cdk5 axis controls the diabetogenic actions of PPARγ. Nature 517:391–395; doi: 10.1038/nature13887.

Becnel LB, Darlington YF, Ochsner SA, Easton-Marks JR Watkins CM, McOwiti A, et al. 2015. Nuclear Receptor Signaling Atlas: Opening Access to the Biology of Nuclear Receptor Signaling Pathways. F.M. Sladek, ed PLOS ONE 10:e0135615; doi:10.1371/journal.pone.0135615.

Bertuloso BD, Podratz PL, Merlo E, de Araújo JFP, Lima LCF, de Miguel EC, et al. 2015. Tributyltin chloride leads to adiposity and impairs metabolic functions in the rat liver and pancreas. Toxicol Lett 235:45–59; doi:10.1016/j.toxlet.2015.03.009.

Calderon-Dominguez M, Sebastián D, Fucho R, Weber M, Mir JF, García-Casarrubios E, et al. 2016. Carnitine Palmitoyltransferase 1 Increases Lipolysis, UCP1 Protein Expression and Mitochondrial Activity in Brown Adipocytes. PloS One 11:e0159399; doi: 10.1371/journal.pone.0159399.

Cappellen D, Luong-Nguyen N-H, Bongiovanni S, Grenet O, Wanke C, Susa M. 2002. Transcriptional program of mouse osteoclast differentiation governed by the macrophage colony-stimulating factor and the ligand for the receptor activator of NFkappa B. J Biol Chem 277:21971–21982; doi: 10.1074/jbc.M200434200.

Chamorro-García R, Sahu M, Abbey RJ, Laude J, Pham N, Blumberg B. 2013. Transgenerational Inheritance of Increased Fat Depot Size, Stem Cell Reprogramming, and Hepatic Steatosis Elicited by Prenatal Exposure to the Obesogen Tributyltin in Mice. Environ Health Perspect 121:359–366; doi:10.1289/ehp.1205701.

Choi JH, Banks AS, Estall JL, Kajimura S, Boström P, Laznik D, et al. 2010. Anti-diabetic drugs inhibit obesity-linked phosphorylation of PPARγ by Cdk5. Nature 466:451–456; doi:10.1038/nature09291.

Choi JH, Banks AS, Kamenecka TM, Busby SA, Chalmers MJ, Kumar N, et al. 2011. Antidiabetic actions of a non-agonist PPARγ ligand blocking Cdk5-mediated phosphorylation. Nature 477:477–481; doi:10.1038/nature10383.

Choi S-S, Kim ES, Koh M, Lee S-J, Lim D, Yang YR, et al. 2014. A novel non-agonist peroxisome proliferator-activated receptor γ (PPARγ) ligand UHC1 blocks PPARγ phosphorylation by cyclin-dependent kinase 5 (CDK5) and improves insulin sensitivity. J Biol Chem 289:26618–26629; doi:10.1074/jbc.M114.566794.

Claussnitzer M, Dankel SN, Kim K-H, Quon G, Meuleman W, Haugen C, et al. 2015. FTO Obesity Variant Circuitry and Adipocyte Browning in Humans. N Engl J Med 373:895–907; doi:10.1056/NEJMoa1502214.

Cohen P, Levy JD, Zhang Y, Frontini A, Kolodin DP, Svensson KJ, et al. 2014. Ablation of PRDM16 and beige adipose causes metabolic dysfunction and a subcutaneous to visceral fat switch. Cell 156:304–316; doi:10.1016/j.cell.2013.12.021.

Cohen P, Spiegelman BM. 2015. Brown and Beige Fat: Molecular Parts of a Thermogenic Machine. Diabetes 64:2346–2351; doi:10.2337/db15-0318.

Feige JN, Auwerx J. 2007. Transcriptional coregulators in the control of energy homeostasis. Trends Cell Biol 17:292–301; doi:10.1016/j.tcb.2007.04.001.

Feige JN, Gelman L, Rossi D, Zoete V, Métivier R, Tudor C, et al. 2007. The endocrine disruptor monoethyl-hexyl-phthalate is a selective peroxisome proliferator-activated receptor gamma modulator that promotes adipogenesis. J Biol Chem 282:19152–19166; doi:10.1074/jbc.M702724200.

Garcia-Vallvé S, Guasch L, Tomas-Hernández S, del Bas JM, Ollendorff V, Arola L, et al. 2015. Peroxisome Proliferator-Activated Receptor γ (PPARγ) and Ligand Choreography: Newcomers Take the Stage. J Med Chem 58:5381–5394; doi:10.1021/jm501155f.

Gburcik V, Cawthorn WP, Nedergaard J, Timmons JA, Cannon B. 2012. An essential role for Tbx15 in the differentiation of brown and “brite” but not white adipocytes. Am J Physiol Endocrinol Metab 303:E1053–1060; doi:10.1152/ajpendo.00104.2012.

Giner XC, Cotnoir-White D, Mader S, Lévesque D. 2015. Selective ligand activity at Nur/retinoid X receptor complexes revealed by dimer-specific bioluminescence resonance energy transfer-based sensors. FASEB J Off Publ Fed Am Soc Exp Biol 29:4256–4267; doi:10.1096/fj.14-259804.

Grün F, Blumberg B. 2006. Environmental obesogens: organotins and endocrine disruption via nuclear receptor signaling. Endocrinology 147:S50–55; doi:10.1210/en.2005-1129.

Grün F, Watanabe H, Zamanian Z, Maeda L, Arima K, Cubacha R, et al. 2006. Endocrine-disrupting organotin compounds are potent inducers of adipogenesis in vertebrates. Mol Endocrinol Baltim Md 20:2141–2155; doi:10.1210/me.2005-0367.

Guan H-P. 2005. Corepressors selectively control the transcriptional activity of PPAR in adipocytes. Genes Dev 19:453–461; doi:10.1101/gad.1263305.

Harms M, Seale P. 2013. Brown and beige fat: development, function and therapeutic potential. Nat Med 19:1252–1263; doi:10.1038/nm.3361.

Hilton C, Karpe F, Pinnick KE. 2015. Role of developmental transcription factors in white, brown and beige adipose tissues. Biochim Biophys Acta 1851:686–696; doi:10.1016/j.bbalip.2015.02.003.

Hu P, Chen X, Whitener RJ, Boder ET, Jones JO, Porollo A, et al. 2013. Effects of parabens on adipocyte differentiation. Toxicol Sci Off J Soc Toxicol 131:56–70; doi:10.1093/toxsci/kfs262.

Imai T, Takakuwa R, Marchand S, Dentz E, Bornert J-M, Messaddeq N, et al. 2004. Peroxisome proliferator-activated receptor gamma is required in mature white and brown adipocytes for their survival in the mouse. Proc Natl Acad Sci U S A 101:4543–4547; doi:10.1073/pnas.0400356101.

Ji X, Chen D, Xu C, Harris SE, Mundy GR, Yoneda T. 2000. Patterns of gene expression associated with BMP-2-induced osteoblast and adipocyte differentiation of mesenchymal progenitor cell 3T3-F442A. J Bone Miner Metab 18: 132–139.

Jornayvaz FR, Shulman GI. 2010. Regulation of mitochondrial biogenesis. Essays Biochem 47:69–84; doi:10.1042/bse0470069.

Juge-Aubry C, Pernin A, Favez T, Burger AG, Wahli W, Meier CA, et al. 1997. DNA binding properties of peroxisome proliferator-activated receptor subtypes on various natural peroxisome proliferator response elements. Importance of the 5’-flanking region. J Biol Chem 272: 25252–25259.

Kajimura S, Seale P, Tomaru T, Erdjument-Bromage H, Cooper MP, Ruas JL, et al. 2008. Regulation of the brown and white fat gene programs through a PRDM16/CtBP transcriptional complex. Genes Dev 22:1397–1409; doi: 10.1101/gad.1666108.

Kanayama T, Kobayashi N, Mamiya S, Nakanishi T, Nishikawa J. 2005. Organotin compounds promote adipocyte differentiation as agonists of the peroxisome proliferator-activated receptor gamma/retinoid X receptor pathway. Mol Pharmacol 67:766–774; doi:10.1124/mol.104.008409.

Kojetin DJ, Matta-Camacho E, Hughes TS, Srinivasan S, Nwachukwu JC, Cavett V, et al. 2015. Structural mechanism for signal transduction in RXR nuclear receptor heterodimers. Nat Commun 6:8013; doi:10.1038/ncomms9013.

Koppen A, Kalkhoven E. 2010. Brown vs white adipocytes: The PPARγ coregulator story. FEBS Lett 584:3250–3259; doi:10.1016/j.febslet.2010.06.035.

Krings A, Rahman S, Huang S, Lu Y, Czernik PJ, Lecka-Czernik B. 2012. Bone marrow fat has brown adipose tissue characteristics, which are attenuated with aging and diabetes. Bone 50:546–552; doi:10.1016/j.bone.2011.06.016.

Lammi J, Perlmann T, Aarnisalo P. 2008. Corepressor interaction differentiates the permissive and non-permissive retinoid X receptor heterodimers. Arch Biochem Biophys 472:105–114; doi:10.1016/j.abb.2008.02.003.

le Maire A, Grimaldi M, Roecklin D, Dagnino S, Vivat-Hannah V, Balaguer P, et al. 2009. Activation of RXR-PPAR heterodimers by organotin environmental endocrine disruptors. EMBO Rep 10:367–373; doi:10.1038/embor.2009.8.

Li X, Pham HT, Janesick AS, Blumberg B. 2012. Triflumizole is an Obesogen in Mice that Acts through Peroxisome Proliferator Activated Receptor Gamma (PPARγ). Environ Health Perspect; doi: 10.1289/ehp.1205383.

Li X, Ycaza J, Blumberg B. 2011. The environmental obesogen tributyltin chloride acts via peroxisome proliferator activated receptor gamma to induce adipogenesis in murine 3T3-L1 preadipocytes. J Steroid Biochem Mol Biol 127:9–15; doi:10.1016/j.jsbmb.2011.03.012.

Lo KA, Sun L. 2013. Turning WAT into BAT: a review on regulators controlling the browning of white adipocytes. Biosci Rep 33; doi:10.1042/BSR20130046.

Michalik L, Zoete V, Krey G, Grosdidier A, Gelman L, Chodanowski P, et al. 2007. Combined Simulation and Mutagenesis Analyses Reveal the Involvement of Key Residues for Peroxisome Proliferator-activated Receptor Helix 12 Dynamic Behavior. J Biol Chem 282:9666–9677; doi: 10.1074/jbc.M610523200.

Murholm M, Dixen K, Qvortrup K, Hansen LHL, Amri E-Z, Madsen L, et al. 2009. Dynamic regulation of genes involved in mitochondrial DNA replication and transcription during mouse brown fat cell differentiation and recruitment. PloS One 4:e8458; doi:10.1371/journal.pone.0008458.

Nolte RT, Wisely GB, Westin S, Cobb JE, Lambert MH, Kurokawa R, et al. 1998. Ligand binding and co-activator assembly of the peroxisome proliferator-activated receptor-gamma. Nature 395:137–143; doi:10.1038/25931.

Oakes ND, Kennedy CJ, Jenkins AB, Laybutt DR, Chisholm DJ, Kraegen EW. 1994. A new antidiabetic agent, BRL 49653, reduces lipid availability and improves insulin action and glucoregulation in the rat. Diabetes 43: 1203–1210.

Pbm de A, A B-L, Mic A-V. 2016. Mitochondrial Actions for Fat Browning and Energy Expenditure in White Adipose Tissue. J Obes Overweight 2; doi:10.15744/2455-7633.2.201.

Pereira-Fernandes A, Vanparys C, Vergauwen L, Knapen D, Jorens PG, Blust R. 2014. Toxicogenomics in the 3T3-L1 cell line, a new approach for screening of obesogenic compounds. Toxicol Sci Off J Soc Toxicol 140:352–363; doi:10.1093/toxsci/kfu092.

Pfaffl MW. 2001. A new mathematical model for relative quantification in real-time RT-PCR. Nucleic Acids Res 29: e45.

Piccolo SR, Withers MR, Francis OE, Bild AH, Johnson WE. 2013. Multiplatform single-sample estimates of transcriptional activation. Proc Natl Acad Sci 110:17778–17783; doi: 10.1073/pnas.1305823110.

Pillai HK, Fang M, Beglov D, Kozakov D, Vajda S, Stapleton HM, et al. 2014. Ligand binding and activation of PPARγ by Firemaster^®^ 550: effects on adipogenesis and osteogenesis in vitro. Environ Health Perspect 122:1225–1232; doi:10.1289/ehp.1408111.

Pittenger MF, Mackay AM, Beck SC, Jaiswal RK, Douglas R, Mosca JD, et al. 1999. Multilineage potential of adult human mesenchymal stem cells. Science 284: 143–147.

Pochetti G, Godio C, Mitro N, Caruso D, Galmozzi A, Scurati S, et al. 2007. Insights into the mechanism of partial agonism: crystal structures of the peroxisome proliferator-activated receptor gamma ligand-binding domain in the complex with two enantiomeric ligands. J Biol Chem 282:17314–17324; doi: 10.1074/jbc.M702316200.

Regnier SM, El-Hashani E, Kamau W, Zhang X, Massad NL, Sargis RM. 2015. Tributyltin differentially promotes development of a phenotypically distinct adipocyte. Obes Silver Spring Md 23:1864–1871; doi: 10.1002/oby.21174.

Ritchie ME, Phipson B, Wu D, Hu Y, Law CW, Shi W, et al. 2015. limma powers differential expression analyses for RNA-sequencing and microarray studies. Nucleic Acids Res 43:e47; doi:10.1093/nar/gkv007.

Riu A, Grimaldi M, le Maire A, Bey G, Phillips K, Boulahtouf A, et al. 2011. Peroxisome proliferator-activated receptor y is a target for halogenated analogs of bisphenol A. Environ Health Perspect 119:1227–1232; doi: 10.1289/ehp.1003328.

Robinson-Rechavi M. 2003. The nuclear receptor superfamily. J Cell Sci 116:585–586; doi:10.1242/jcs.00247.

Rosell M, Kaforou M, Frontini A, Okolo A, Chan Y-W, Nikolopoulou E, et al. 2014. Brown and white adipose tissues: intrinsic differences in gene expression and response to cold exposure in mice. Am J Physiol Endocrinol Metab 306:E945–964; doi:10.1152/ajpendo.00473.2013.

Saeed H, Iqtedar M. 2015. Aberrant gene expression profiles, during in vitro osteoblast differentiation, of telomerase deficient mouse bone marrow stromal stem cells (mBMSCs). J Biomed Sci 22:11; doi:10.1186/s12929-015-0116-4.

Schilling AF, Schinke T, Münch C, Gebauer M, Niemeier A, Priemel M, et al. 2005. Increased bone formation in mice lacking apolipoprotein E. J Bone Miner Res Off J Am Soc Bone Miner Res 20:274–282; doi:10.1359/JBMR.041101.

Schulman IG, Shao G, Heyman RA. 1998. Transactivation by retinoid X receptor-peroxisome proliferator-activated receptor gamma (PPARgamma) heterodimers: intermolecular synergy requires only the PPARgamma hormone-dependent activation function. Mol Cell Biol 18: 3483–3494.

Seale P. 2015. Transcriptional Regulatory Circuits Controlling Brown Fat Development and Activation. Diabetes 64:2369–2375; doi:10.2337/db15-0203.

Shiizaki K, Yoshikawa T, Takada E, Hirose S, Ito-Harashima S, Kawanishi M, et al. 2014. Development of yeast reporter assay for screening specific ligands of retinoic acid and retinoid X receptor subtypes. J Pharmacol Toxicol Methods 69:245–252; doi:10.1016/j.vascn.2014.01.007.

Sidossis L, Kajimura S. 2015. Brown and beige fat in humans: thermogenic adipocytes that control energy and glucose homeostasis. J Clin Invest 125:478–486; doi:10.1172/JCI78362.

Soccio RE, Chen ER, Lazar MA. 2014. Thiazolidinediones and the promise of insulin sensitization in type 2 diabetes. Cell Metab 20:573–591; doi:10.1016/j.cmet.2014.08.005.

Spoto B, Di Betta E, Mattace-Raso F, Sijbrands E, Vilardi A, Parlongo RM, et al. 2014. Pro- and antiinflammatory cytokine gene expression in subcutaneous and visceral fat in severe obesity. Nutr Metab Cardiovasc Dis NMCD 24:1137–1143; doi:10.1016/j.numecd.2014.04.017.

Stinkens R, Goossens GH, Jocken JWE, Blaak EE. 2015. Targeting fatty acid metabolism to improve glucose metabolism. Obes Rev Off J Int Assoc Study Obes 16:715–757; doi:10.1111/obr.12298.

Subramanian A, Tamayo P, Mootha VK, Mukherjee S, Ebert BL, Gillette MA, et al. 2005. Gene set enrichment analysis: a knowledge-based approach for interpreting genome-wide expression profiles. Proc Natl Acad Sci U S A 102:15545–15550; doi:10.1073/pnas.0506580102.

Timmons JA, Pedersen BK. 2009. The importance of brown adipose tissue. N Engl J Med 361:415–416; author reply 418-421; doi:10.1056/NEJMc091009.

Tontonoz P, Hu E, Spiegelman BM. 1994. Stimulation of adipogenesis in fibroblasts by PPAR gamma 2, a lipid-activated transcription factor. Cell 79: 1147–1156.

Tsukamoto Y, Ishihara Y, Miyagawa-Tomita S, Hagiwara H. 2004. Inhibition of ossification in vivo and differentiation of osteoblasts in vitro by tributyltin. Biochem Pharmacol 68:739–746; doi:10.1016/j.bcp.2004.04.020.

Tudor C, Feige JN, Pingali H, Lohray VB, Wahli W, Desvergne B, et al. 2007. Association with Coregulators Is the Major Determinant Governing Peroxisome Proliferator-activated Receptor Mobility in Living Cells. J Biol Chem 282:4417–4426; doi:10.1074/jbc.M608172200.

Wang H, Liu L, Lin JZ, Aprahamian TR, Farmer SR. 2016. Browning of White Adipose Tissue with Roscovitine Induces a Distinct Population of UCP1(+) Adipocytes. Cell Metab 24:835–847; doi: 10.1016/j.cmet.2016.10.005.

Wang P, Renes J, Bouwman F, Bunschoten A, Mariman E, Keijer J. 2007. Absence of an adipogenic effect of Rosiglitazone on mature 3T3-L1 adipocytes: increase of lipid catabolism and reduction of adipokine expression. Diabetologia 50:654–665; doi:10.1007/s00125-006-0565-0.

Watt J, Schlezinger JJ. 2015. Structurally-diverse, PPARγ-activating environmental toxicants induce adipogenesis and suppress osteogenesis in bone marrow mesenchymal stromal cells. Toxicology 331:66–77; doi:10.1016/j.tox.2015.03.006.

Wensveen FM, Valentić S, Šestan M, Turk Wensveen T, Polić B. 2015. The “Big Bang” in obese fat: Events initiating obesity-induced adipose tissue inflammation. Eur J Immunol 45:2446–2456; doi: 10.1002/eji.201545502.

Xiao Y, Cui J, Li Y-X, Shi Y-H, Le G-W. 2010. Expression of genes associated with bone resorption is increased and bone formation is decreased in mice fed a high-fat diet. Lipids 45:345–355; doi: 10.1007/s11745-010-3397-0.

Xiao Y, Cui J, Shi Y, Le G. 2011. Lipoic acid increases the expression of genes involved in bone formation in mice fed a high-fat diet. Nutr Res N Y N 31:309–317; doi: 10.1016/j.nutres.2011.03.013.

Yanik SC, Baker AH, Mann KK, Schlezinger JJ. 2011. Organotins are potent activators of PPARγ and adipocyte differentiation in bone marrow multipotent mesenchymal stromal cells. Toxicol Sci Off J Soc Toxicol 122:476–488; doi:10.1093/toxsci/kfr140.

Yao W, Cheng Z, Busse C, Pham A, Nakamura MC, Lane NE. 2008. Glucocorticoid excess in mice results in early activation of osteoclastogenesis and adipogenesis and prolonged suppression of osteogenesis: a longitudinal study of gene expression in bone tissue from glucocorticoid-treated mice. Arthritis Rheum 58:1674–1686; doi:10.1002/art.23454.

Zuo Z, Chen S, Wu T, Zhang J, Su Y, Chen Y, et al. 2011. Tributyltin causes obesity and hepatic steatosis in male mice. Environ Toxicol 26:79–85; doi: 10.1002/tox.20531.

